# WUREN: Whole-modal fUsion Representation for protein interaction interfacE predictioN

**DOI:** 10.1101/2022.12.22.521634

**Authors:** Xiaodong Wang, Xiangrui Gao, Xuezhe Fan, Zhe Huai, Mengcheng Yao, Tianyuan Wang, Xiaolu Huang, Lipeng Lai

**Affiliations:** Xtalpi Innovation Center, Beijing, China; Xtalpi Investment, Suzhou, China

## Abstract

Proteins are one of the most important components in life, and the research on protein complex and the development of protein or antibody drugs relies on effective representation of proteins. Both experimental methods like cryo-electron microscopy and computational methods like molecular dynamic simulation suffer from high cost, long time investment and low throughput, and cannot be used in large-scale studies. Some examples of artificial intelligence for protein complex prediction tasks show that different representations of proteins have their own limitations. This paper constructs a multimodal model named WUREN (Whole-modal fUsion Representation for protein interaction interfacE predictioN), which effectively fuses sequence, graph, and structural features. WUREN has achieved state-of-the-art performance on both the antigen epitope prediction task and the protein-protein interaction interface prediction task, with AUC-PR reaching 0.462 and 0.516, respectively. Our results show that WUREN is a general and effective feature extraction model for protein complex, which can be used in the development of protein-based drugs. Furthermore, the general framework in WUREN can be potentially applied to model similar biologics to proteins, such as DNA and RNA.

## Introduction

Proteins are ubiquitous in life and biological processes. Deeper understanding of the relations between the composition of specific proteins/protein complexes will help us better understand the function of life itself, combat different diseases etc. For example, a better cognition of how B Cell Receptors (BCRs) recognize antigens or how and where antibodies bind to antigens can largely promote our capabilities in discovering and developing new antibodies, vaccines, or other therapies for other immune-related diseases.

Many technologies are used for in-depth study of proteins, from experimental and computational perspectives. For instance, to determine the relationship between the characterization and function of protein complex, the structure of the protein was obtained experimentally by cryo-electron microscopy (Cryo-EM), X-ray, and nuclear magnetic resonance (NMR) ^1,2,3^. However, these methods have their own limitations. Cryo-EM takes an average of 2 weeks and $50,000 to obtain a complex with a resolution of 1.5 Å. X-ray needs to obtain protein crystallization first. NMR can only be used for structure elucidation of small complexes with equivalent weights less than 4OkDa. In addition to the above shortcomings, all these methods are low-throughput methods and cannot be used in large-scale application scenarios such as protein and antibody drug development.

In computations, researchers usually use molecular dynamic simulation methods to study the structure and function of proteins. SnugDock is one of the tools used to study the molecular dynamic process of protein complex interactions^4^. But such methods usually consume an amount of computing resources and time and cannot achieve large-scale applications.

Recent progresses in artificial intelligence and deep learning shed light on an alternative approach to help us understand proteins^5,6^. AlphaFold2 shows us a successful example of how much information we can extract from the protein sequence merely^5^. Following work like AF2-Multimer and xTrimo-Multimer^6^, expands the application of protein structure prediction to the scenario of protein complex/multimers. As protein-protein interaction (PPI) interface prediction task being one of the key challenges in understanding the protein functions, various other methods based on artificial intelligence are published in this direction^7,8,9^. MaSIF predicts PPI through fusion of chemical and point cloud structure features^7^. PECAN predicts antigen-antibody binding surface through Graph features^8^. PINet predicts antigen-antibody binding surfaces by fusing physicochemical features and point cloud structural features^9^. All these examples show that different features, like sequence features, graph features and structure features, are effective in characterizing proteins. One inevitable question to ask is whether these features encode the same information, or when they work together can improve the performance. Theoretically, the sequences of proteins encode all the information to determine the structures, dynamics, and functions of the proteins. However, when we confront limited training data, auxiliary information may be useful to help the learning process and hence improve the model performance. As shown in previous protein structure prediction methods AlphaFold2, DCA-fold and RaptorX^5,10,11^, when evolutionary information is considered through Multiple Sequence Alignment (MSA), the prediction accuracy is increased. In this study, we show that graphic/ topological (graph) and spatial (structure) information complement with sequential features, and when they are merged fully or partially, the performance of our models is improved in general in relevant tasks.

Achieving effective multimodal feature fusion is very challenging. The difficulty lies in the need to be able to judge the confidence and correlation of each modality, and to achieve multimodal feature alignment and data registration.

In this paper, we build a method called WUREN (Whole-modal fUsion Representation for protein interaction interfacE predictioN), named after a famous Chinese desert, made by a mix of various ingredients. As shown in Figure 1.b, without the loss of generality, Transformer^12^, Graph Convolution Neural Network (GCN) and PointNet++^13,14^, are applied in WUREN to extract sequence, graph, and structure information respectively. Operationally, these methods can be replaced by other same function methods like PointNet, Graph Attention Network (GAT) etc.^15,16,17,18^, within the framework of WUREN according to the need of downstream tasks.

**Fig. 1.**
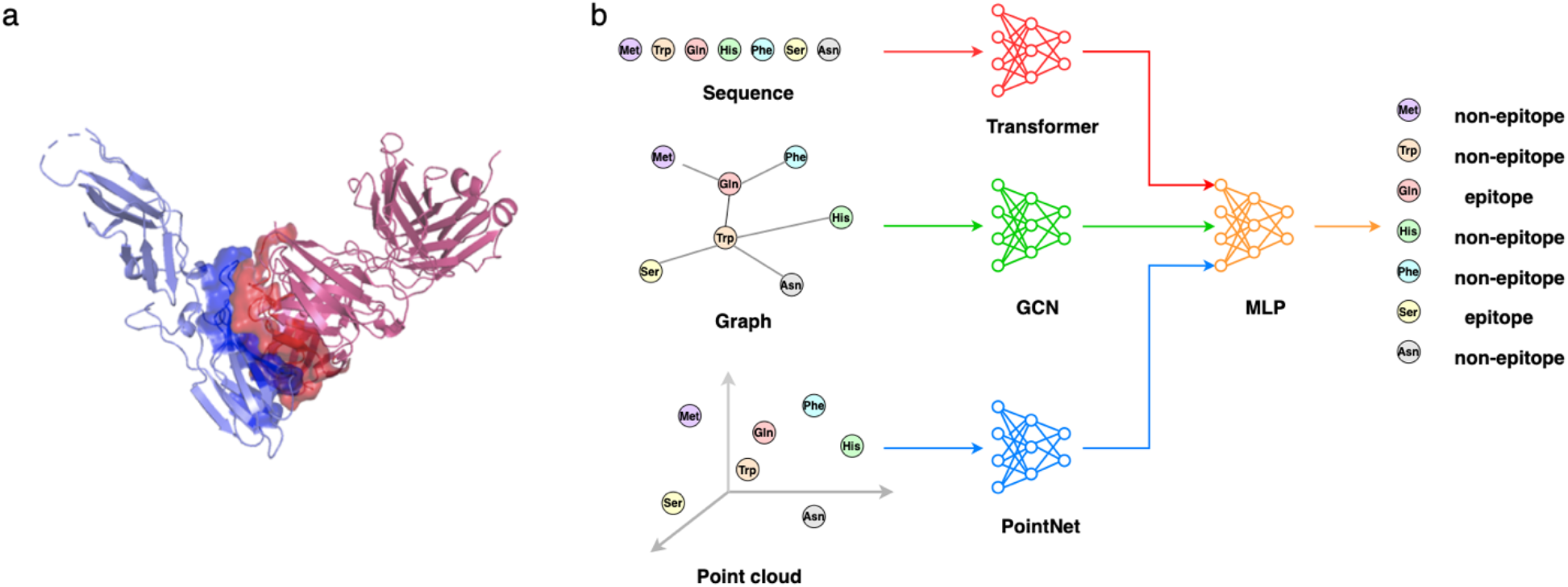
Schematic diagram of epitope prediction process. **a,** schematic diagram of the inter-surface of antigen-antibody complex, including the epitope, the region on the antigen that the antibody specifically bin. Example diagram of a complex (PDB ID: 1ahw) between IMMUNOGLOBULIN (blue) and the antibody (red). **b,** the schematic diagram of WUREN model framework. WUREN is composed of transformer, GCN and PointNet, which realizes the effective fusion of sequential features, topological features, and structural features.

WUREN achieves a state-of-the-art result AUC-PR of 0.462 on the epitope prediction benchmark EpiPred and outperforms the multimer methods on newly released SAbDab dataset^19,20^. Figure 1.a shows a data example of an antigen-antibody complex. Beyond the epitope prediction task, on the PPI prediction benchmark MaSIF^7^, the model again achieves state-of-the-art performance with AUC-PR of 0.512. It is proved by ablation experiments that features from each dimension in the model play an important role.

Our results show that the multimodal model WUREN proposed in this paper is a general and effective model for protein complex representation, which can be used in protein complex-related research and in the development of protein-based drugs. Furthermore, the general idea/framework in WUREN can be potentially applied to model similar biologics to proteins, such as DNA, RNA etc.

## Result

### WUREN

We construct a deep learning model as shown in Figure 2.a, which consists of four modules, Cross Attention PointNet++ (CAP), Cross Attention GCN (CAG), Cross Attention Transformer (CAT), and Multilayer Perceptron (MLP).

**Fig. 2.**
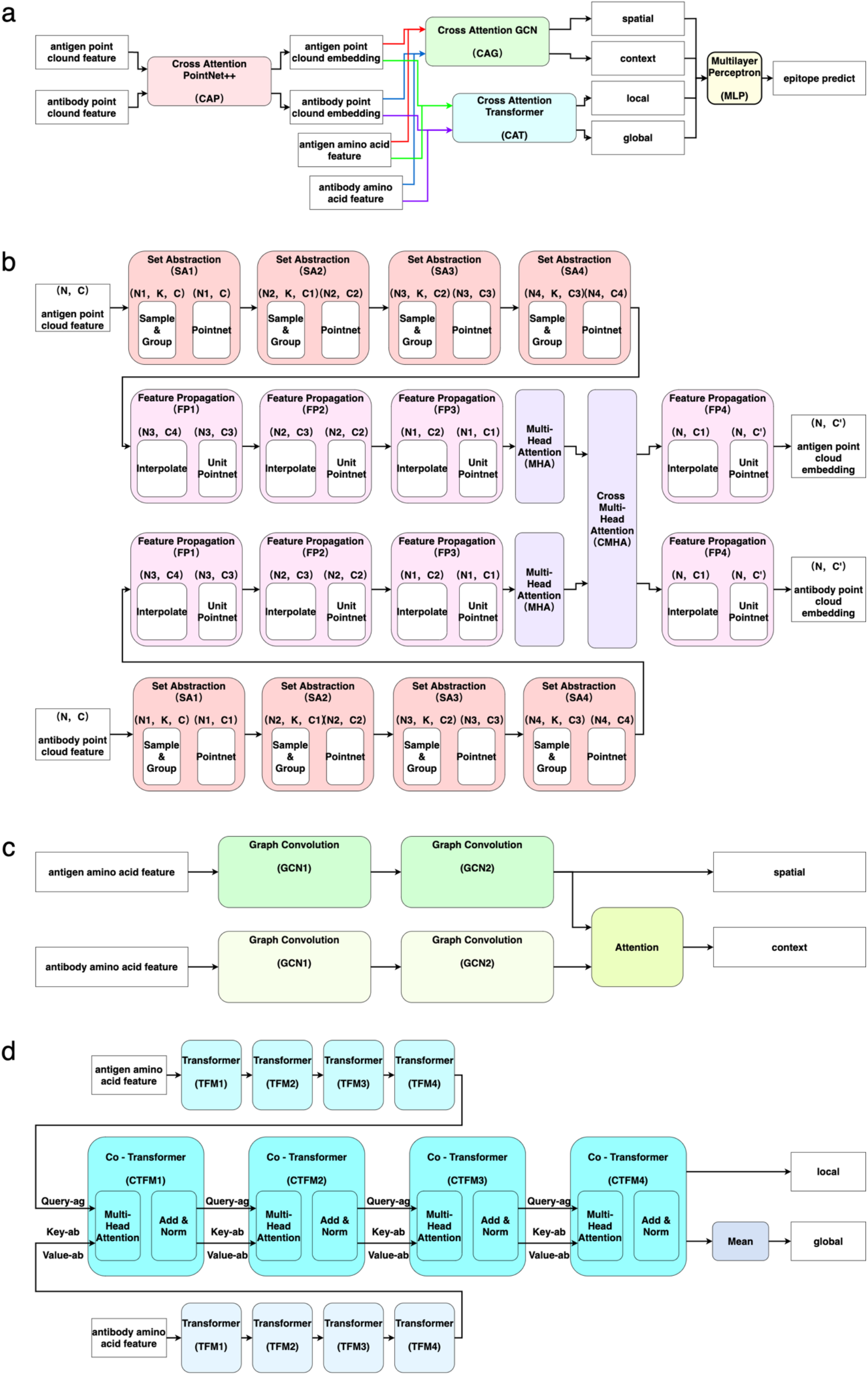
WUREN. **a,** Overall framework of the WUREN model. **b,** Diagram of Cross Attention PointNet++ (CAP) block. **c,** Diagram of Cross Attention GCN (CAG) block. **d,** Diagram of Cross Attention Transformer (CAT) block.

The structure of the CAP module is shown in Figure 2.b. This module is mainly composed of two PointNet++ models, two Self Multi-Head Attention layers and one Cross Multi-Head Attention layer. The PointNet++ model consists of 4 Set Abstraction (SA) sampling modules and 4 Feature Propagation (FP) upsample modules. The point cloud features of antigens and antibodies are extracted into point cloud embedding by the CAP module.

The structure of the CAG module is shown in Figure 2.c. The amino acid features of the antigen and antibody in this module are extracted by two GCN layers, respectively. The graph feature of the antigen itself is spatial, and the context information is obtained after the cross attention of the antigen and antibody is calculated.

The structure of the CAT module is shown in Figure 2.d. The amino acid features of antigens and antibodies in this module are extracted through 4 layers of Transform to extract the embedding, and then the embeddings of the two are input into four continuously Co-Transformer layers to extract Cross Multi-Head Attention^21^. In the Transformer module, in addition to using the absolute position information of each amino acid, we also added the T5 relative position information which representing the distance between every two amino acids^22^. In the Co-Transformer layer, the features of the antigen are used as Query, and the features of the antibody are used as Key and Value, and the attention weight between each two amino acids of the antigen and the antibody is calculated. After processing by this module, we obtained the local information of each amino acid and the global information after averaging all amino acid features.

With the model framework described above, to synergically fuse spatial, topological, and sequential information, the following training steps are performed:

Firstly, we train the CAP model using the features of the antigen and antibody point clouds. Secondly, with the CAP model parameters fixed, we obtain the point cloud embedding of antigens and antibodies through the model and aggregate the point cloud embedding into amino acid level embedding. Thirdly, we concatenate the embedding obtained by the point cloud model with other amino acid level features and train the CAG and CAT models as a result, the spatial, context, local and global information of each amino acid is obtained. Finally, the spatial, context, local and global features are concatenated, and the MLP prediction model is trained to obtain the prediction probability of whether each amino acid is in the epitope set.

After the above process, the spatial information extracted by the point cloud CAP module, the topological information extracted by the graph CAP module, and the sequence information extracted by the sequence CAT module are effectively fused to achieve an effective amino acid representation.

### Epitope prediction performance on EpiPred

We use 108 training sets and 30 test sets consistent with the EpiPred paper^19^. To avoid model overfitting, we use the 44 complexes in the BM dataset that are not in the training set as the validation set^23^. To ensure that the point cloud data of antigens and antibodies meet the input requirements of the model, we sample the antigen and antibody point clouds using the Farthest Point Sampling (FPS) method and obtained 8192 point cloud samples respectively. This process improves the training efficiency of point cloud model, without sacrificing model’s performance. To make the point cloud model more robust, we perform data augmentation on the point cloud data, and each data is subjected to random translation and rotation transformations. Finally, we normalize both the point cloud and amino acid features to avoid the worsening of the model performance due to the singular values of some features.

On the EpiPred dataset, we compare the performance of WUREN with representative models EpiPred, PECAN and PINet^19,8,9^. Area under the precision recall curve (AUC-PR), area under the receiver operating characteristics curve (AUC-ROC), precision and recall are used as test metrics, and the corresponding functions in the sklearn metric module are used for calculating these metrics^24^.

As shown in Figure 3.a, on the 30 test data of EpiPred, WUREN realizes significant improvement in AUC-PR (0.462), AUC-ROC (0.877) and precision (0.359) compared with EpiPred, PECAN and PINet, and achieves the results of current state-of-the-art. Figure 3.d is the scatter plots of the penultimate layer of complex ‘4etq’ embedding after PCA. The positive and negative samples are randomly distributed in the embedding of PINet, but they gathers to specific clusters in the embedding of WUREN.

**Fig. 3.**
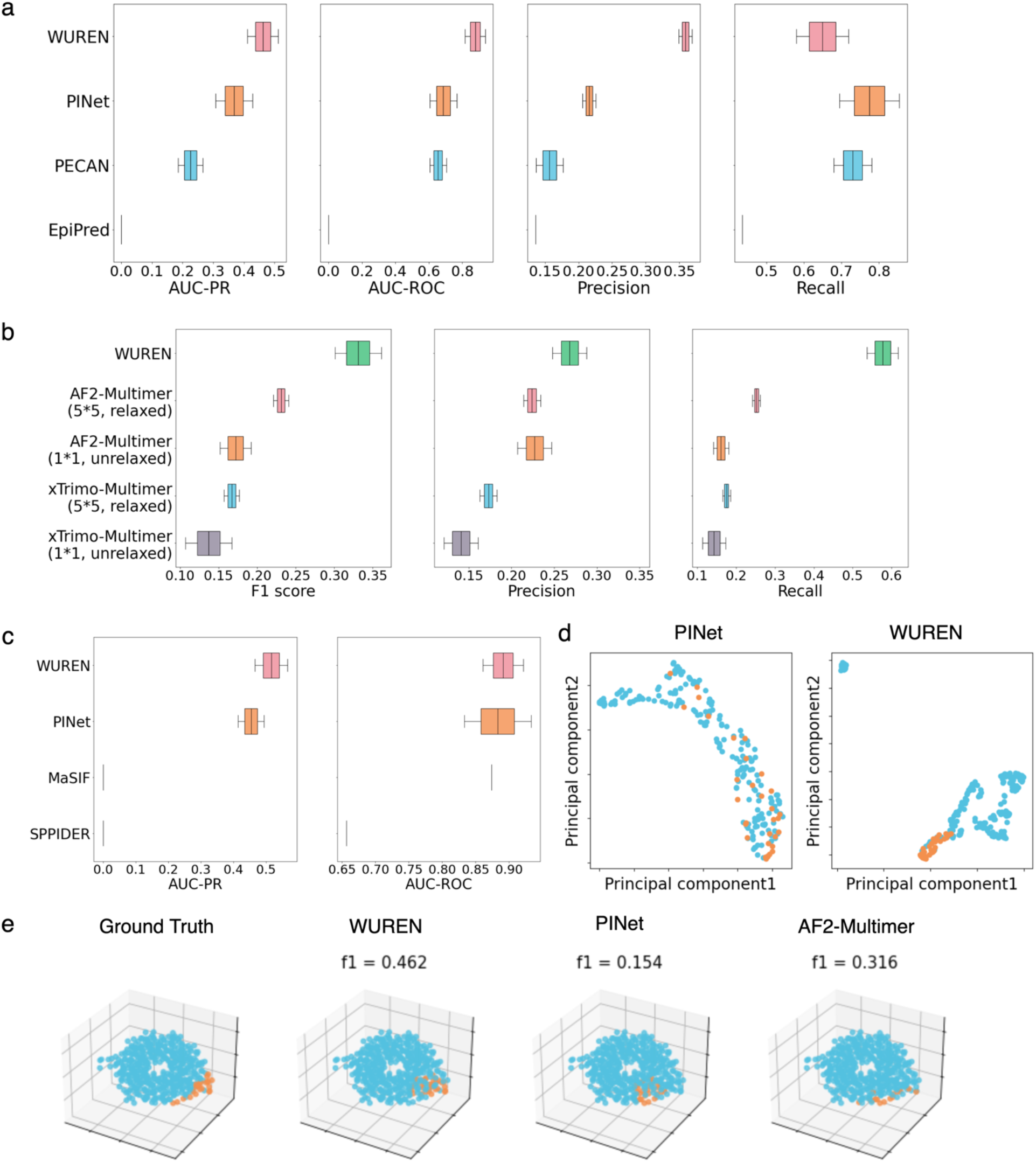
Results on EpiPred, SAbDab and MaSIF benchmark. **a,** WUREN gets significant improvement in AUC-PR (0.462), AUC-ROC (0.877) and precision (0.359) on EpiPred benchmark and achieves the results of current state-of-the-art. **b,** WUREN reaches the best results on SAbDab’s newly released datasets in general. **c,** WUREN attains the best performance on MaSIF benchmark, with AUC-PR 0.516 and AUC-ROC 0.891. **d,** the scatter plots of the penultimate layer’s embedding after PCA, orange represents positive samples, blue represents negative samples. **e,** 3D scatters plots of prediction results of different models, orange represents positive samples, blue represents negative samples.

### Epitope prediction performance on SAbDab

We collect data of 1000 complexes newly released by SAbDab after 2021^20^. We compare them with our training set and eliminate sequences with a similarity identity greater than 50%. As a result, we finally obtain 77 complex data, which is used as the test dataset to compare our model with protein complex structure prediction models, AF2-Multimer and xTrimo-Multimer. F1 score, precision and recall are used as test metrics.

Operationally, two setups of AF2-Multimer and xTrimo-Multimer are used for comparison. One uses a single model to predict a single structure. The other utilizes 5 models, each predicting 5 structures. In the latter configuration, the predicted structure is the best of the 25 structures after relaxation, picked according to the prediction confidence. After obtaining the predicted complex structure, we calculated the pairwise amino acid distance between the antigen and the antibody to obtain the predicted epitope with a threshold of 4.5 Å.

As shown in Figure 3.b, WUREN implements the best results on this dataset in general. Figure 3.e shows the prediction results of different models using the complex ‘3t3p’ as an example.

In addition, AF2-Multimer outperforms xTrimo-Multimer in corresponding settings. It also shows that, for xTrimo-Multimer and AF2-Multimer, the results from multi-model settings with structure relaxation are better than those of single model settings without structure relaxation.

### PPI prediction performance on MaSIF

P. Gainza et al. integrated the PRISM dataset^7,25^, ZDock dataset^26^, PDBBind dataset and SAbDab dataset^27,20^, and filtered out the sequence similarity identity greater than 30% by using psi-cd-hit^28^. In the end, a dataset of 3003 training data and 357 testing data is constructed. We use the same dataset for training and testing to validate our method for other protein-related tasks like PPI prediction. We conduct PPI prediction experiments on the MaSIF dataset, and compare our results with SPPIDER^29^, MaSIF and PINet. To make a parallel comparison with the data in the reference, we use area under the precision recall curve (AUC-PR) and area under the receiver operating characteristics curve (AUC-ROC) as the metric function.

As shown in Figure 3.c, our method WUREN achieves the best performance, with AUC-PR 0.516 and AUC-ROC 0.891. This indicates the potential of WUREN to be generalized to other protein related tasks and broader applications of WUREN to other similar scenarios, e.g., modeling of DNA, RNA, etc.

### Model ablation experiment results

An ablation test is performed to analyze the contribution of each of the three modules that extract sequential, topological, and spatial information respectively. As shown in Figure 4.a, Figure 4.b, and Figure 4.c, on EpiPred’s 30 test data, we conduct ablation experiments of the WUREN model. Overall, the performance is improved as the complexity of the model increases when more representations are fused together.

**Fig. 4.**
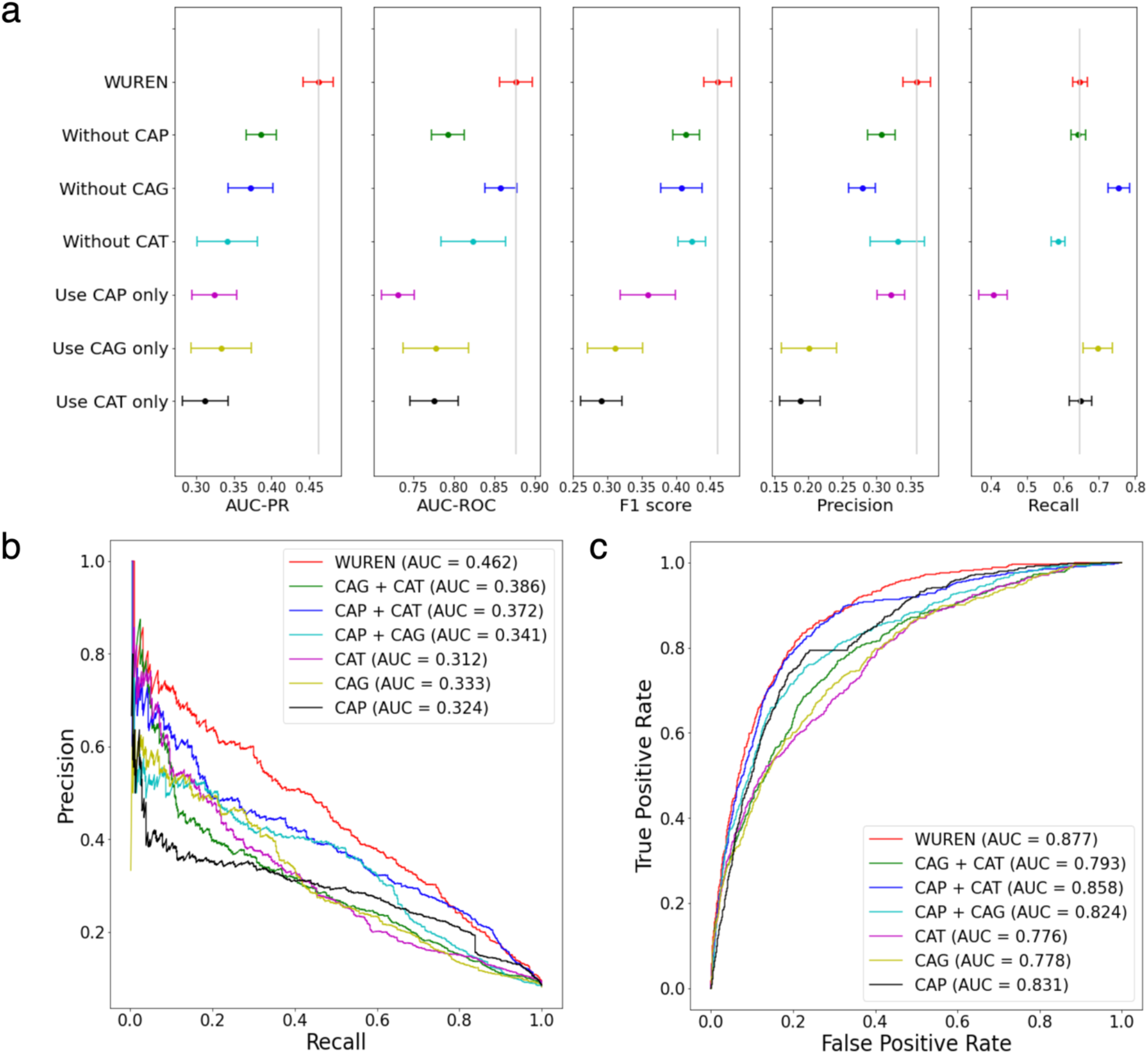
Results of model ablation test. **a,** Ablation results on EpiPred benchmark, the AUC-PR, AUC-ROC, f1 score and precision of WUREN decreases significantly with the reduction of the constituent modules. **b,** AUC-PR is improved as the complexity of the model increases when more representations are merged. **c,** AUC-ROC is improved as the complexity of the model increases when more representations are merged.

### Contributions of attention

Taking the complex ‘4g3y’ as an example, the attention weight matrix of the single CAG module is shown as Figure 5.a, and the attention weight matrix of WUREN with all three modules included (full model) is shown as Figure 5.b. The rows and columns in the figure represent antigens and antibodies, respectively, with black/gray color representing amino acids on non-binding surfaces and green/blue color representing amino acids on paratope and epitope. As shown in these two figures, both the single CAG module and WUREN pay more attention to amino acids on the epitope than those on the non-binding surface, indicating that both models have learnt the relevant features to distinguish the binding and non-binding surfaces. Furthermore, as indicated in the right figure, WUREN, the full model, pays more attention to the location of the binding surface than the model using only CAG, indicating that fusion of sequential, topological, and spatial features captures features more relevant to the binding surface. The results of the attention matrix analysis are consistent with the performance improvement and the ablation test of WUREN (Figure 4).

**Fig. 5.**
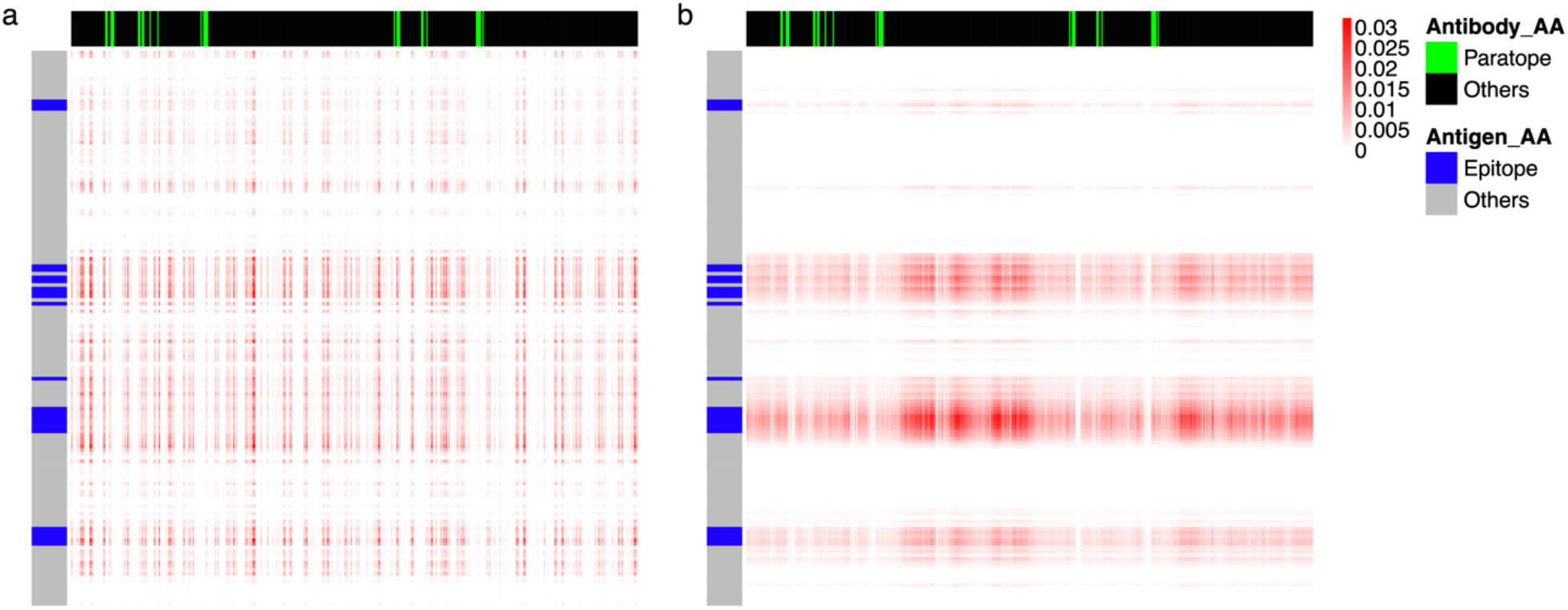
Attention weight map. **a,** The attention weight map of the single CAG module. **b,** The attention weight map of WUREN model.

## Discussion

Besides the main results shown here, we notice in our study that selecting and adding more suitable amino acid features significantly improves model performance. In the case where only the CAG module is used for modeling, when the dimension of amino acid representation is increased from 63 as in PECAN to 136 as in this study, the AUC-PR increases from 0.226 to 0.338. The 136 features are selected from 183 commonly used amino acid features, such as amino acid type, absolute solvent accessible surface area etc., by random forest algorithm, which contribute more than others to the epitope prediction task. The same procedure can be applied to improve models for other similar tasks.

In addition, we found that post-processing of the results also improves the accuracy of epitope predictions. By clustering the predicted epitopes, selecting the clusters’ central points, and performing nearest neighbor sampling according to the central points, the F1 score on the test set can be improved by approximately 10%.

Furthermore, as one may expect, adding more data improves the performance of the model. We collect all SAbDab data and exclude the data whose maximum sequence similarity with 30 test samples is greater than 50%. The remaining 1075 data is used for model training, and the AUC-PR of the resulting model on the test set is improved to 0.574.

## Conclusion

Proteins are part of the foundation of life. The diverse functions of proteins are tightly related to their structures, which are largely determined by the unique sequences of each protein and the interactions between amino acids. Computational approaches provide an efficient way to understand the function of proteins from the compositions. Although we can decipher lots of information just from the sequences of amino acids or just from the structure, considering the limitation of data we can learn from, a practical model should make better use of the data by utilizing suitable features from different representations.

In this paper, a multimodal protein complex characterization framework WUREN is proposed, which fully integrates one-dimensional sequence features, two-dimensional graph features and three-dimensional structural features. To demonstrate the advantage of WUREN, we realize the model with three specific modules, CAT, CAG and CAP, to extract sequence features, graph features and structural features, respectively. In epitope prediction task, our model achieves SOTA results on AUC-PR, AUC-ROC, and precision on EpiPred dataset and better performance on 77 newly released data of SAbDab compared to some of the advanced Multimer methods. Ablation experiments showed that sequence features, graph features and structural features all play important roles. In addition, by comparing the attention maps of a single CAG module and WUREN, it shows that the latter is more effective in learning features that are more relevant to the epitope prediction task.

By testing WUREN on the PPI prediction task with dataset MaSIF, we also demonstrated that the framework from WUREN can be generalized to other protein related tasks. We believe modeling of other modalities, like DNA, RNA, etc., which share similar linear sequential structure as proteins, may also benefit from the framework of WUREN.

There are still some directions to improve the model. Firstly, since the calculation process of aggregating point cloud embedding into amino acid embedding is non-differentiable, the model in this paper needs to be trained in two stages, which increases the training difficulty and may also reduce the overall performance of the model. Secondly, the model requires the knowledge of the structure of proteins under investigation, which can be difficult to obtain in some scenarios. Further application of WUREN to those cases when structure information is not available will rely on the continuous improvement of protein structure prediction models, especially for models predicting antibody structures.

In conclusion, here we propose a new multimodal deep learning model WUREN for tasks related to proteins or protein complexes. The model is shown to achieve SOTA performances in example tasks and can be potentially generalized to different protein related tasks or other biologics. Further investigation of this model will help us gain a better understanding of proteins and other biologics, and potentially aid the design of new therapies.

## Methods

### Problem statement

We select the task of epitope prediction as the example to demonstrate the performance of WUREN. The goal is to predict the region of the antigen surface bound to the antibody, in amino acid units. For data process, we use a variety of algorithms and tools to extract physical, chemical, and structural features of antigens and antibodies in units of amino acids and point clouds, respectively. To calculate the labels, we first calculate the distance between each point cloud of the antigen and the antibody and mark the point cloud with a distance less than 2 Å as 1, indicating that the point cloud belongs to the binding surface, and the other point clouds are marked as 0. Similarly, we calculate the distance between each amino acid of the antigen and the antibody. The amino acid with a distance less than 4.5 Å is marked as 1, indicating that the amino acid belongs to the epitope region, and the remaining amino acids are marked as 0. Finally, we use the point cloud data to train the deep learning model, then use 2 Å as the threshold to gather the obtained point cloud embedding into amino acid features, combine the prepared amino acid features to train the final model, and predict the probability that each amino acid belongs to the epitope region, to complete the epitope prediction.

### Features

We extract point clouds and amino acid features of antigens and antibodies.

#### Point cloud features

We first process the PDB file using PDB2PQR^30,31^, removing solvent molecules and filling in missing atoms. Then we extract the surface meshes of antigens and antibodies to obtain point cloud data. We process the point cloud data using APBS to obtain Poisson–Boltzmann electrostatics for each point cloud^32^. For this part, we obtain the point cloud features of antigen and antibody respectively. Each point cloud contains 3-dimensional spatial coordinate Gi information (x(i), y(i), z(i)) and 11-dimensional charge Ei information.

#### Amino acid features

We use a variety of algorithms and tools for the extraction of antigen and antibody amino acid features. (1) One-hot encoding: we perform one-hot encoding for all amino acids with all uncommon amino acids are classified into one class, and finally a feature with a length of 21 is obtained for each amino acid. (2) Neighbor composition: we calculate the frequency of 20 common amino acids within its 8 Å distance for each amino acid. This provides a feature of length 20. (3) Absolute and relative solvent accessible surface area: we use PyRosetta^33^, which encapsulated by the powerful platform Rosseta^34^, to calculate the absolute and relative solvent accessible surface area of each amino acid, forming a feature of length 2. (4) Position-Specific Scoring Matrix (PSSM): we use PSI-BLAST to calculate the PSSM of each amino acid to obtain a feature of length 20^35^. (5) Peptides: we use R Package Peptides to extract the features of amino acids^36^, and obtain physicochemical features such as Cruciani Properties^37^, forming a feature of length 66. (6) Others: we calculate residue depth, residue adjacency degree, average B-factor, Isoelectric and molecular weight, forming a feature of length 6. Finally, we compute a feature of length 136 for each amino acid of the antigen and antibody.

### Implementation Details

We use a single A100 GPU with 15G memory to train the model.

#### Point cloud CAP model training

Adam optimizer is used with learning rate 0.0001 and weight decay 0.001^38^. We use Cross Entropy as the loss function^39^, and the weight is set to 0.075 to reduce the impact of the imbalance of positive and negative samples on model training. Batch size is set to 5, which means that 5 complete antigen and antibody point cloud pairs are processed each time. The dropout rate is set to 0.6. The training epoch is 300.

#### Splice point cloud embedding and amino acid features

We fix the parameters of the point cloud CAP model and extract the antigen and antibody point cloud embeddings with a length of 128 through the CAP model. We correspond the coordinates of the point cloud with the coordinates of the amino acids in the PDB file, extract all point clouds with less than 2 Å for each amino acid, and average the embedding of these point clouds to obtain an embedding with a length of 128 for each amino acid. As a result, the point cloud embedding is converted into amino acid embedding. We splice the amino acid embedding of length 128 containing spatial structure information obtained by the CAP module and the amino acid feature of length 136 obtained in the data preparation stage and obtained a feature of length 264 for each amino acid.

#### CAG and CAT model training

We use Adam as the optimizer with learning rate 0.0001 and weight decay 0.001. We use Cross Entropy as the loss function, and the weight is set to 0.04 to reduce the impact of the imbalance of positive and negative samples on model training. Batch size is set to 32, which means 32 amino acids of the antigen are processed each time. The dropout rate is set to 0.6. The training epoch is 100.

## Supporting information

Supplemental Files for WUREN

## Data availability

The data used in this article will be provided after the paper is accepted ‘in principle’.

## Code availability

The WUREN code was implemented in Python using the deep learning framework of PyTorch. Code, trained models, and scripts reproducing the experiments of this article will be provided after the paper is accepted ‘in principle’.

## Acknowledgements

The authors thank Chaohui Gong for helpful comments, and Genwei Zhang, Ruyu Wang, Lianjun Zheng for proofreading.

## Author contributions

X.W and L.L conceived this research. X.F, Z.H, M.Y and T.W curated the dataset. X.W performed data analysis. X.W and X.G devised deep learning algorithms. X.W conducted the experiments. X.W and L.L wrote and modified the paper. X.H and L.L supervised this work.

## Additional information

**Supplementary information**

**Correspondence and requests for materials** should be addressed to Lipeng Lai

**Peer review information**

**Reprints and permissions information**

**Publisher’s note**

