## Supplemental Files for WUREN for "WUREN: Whole-modal fUsion Representation for protein interaction interfacE predictioN"

Supplementary Information

### Performance on EpiPred Dataset

WUREN achieves significant improvement in AUC-PR (0.462), AUC-ROC (0.877) and Precision (0.359) compared with EpiPred, PECAN and PINet, and achieves the results of current state-of-the-art.

Supplementary Table. 1: Test result table of WUREN on EpiPred datasets

| Methods | AUC-PR | AUC-ROC | Precision | Recall |
| --- | --- | --- | --- | --- |
| Epipred | Na | Na | 0.136 | 0.436 |
| PECAN | 0.226 ± 0.04 | 0.655 ± 0.05 | 0.157 ± 0.02 | 0.730 ± 0.05 |
| PINet | 0.368 ± 0.06 | 0.687 ± 0.08 | 0.216 ± 0.01 | **0.774 ± 0.08** |
| WUREN | **0.462 ± 0.05** | **0.877 ± 0.06** | **0.359 ± 0.01** | 0.647 ± 0.07 |

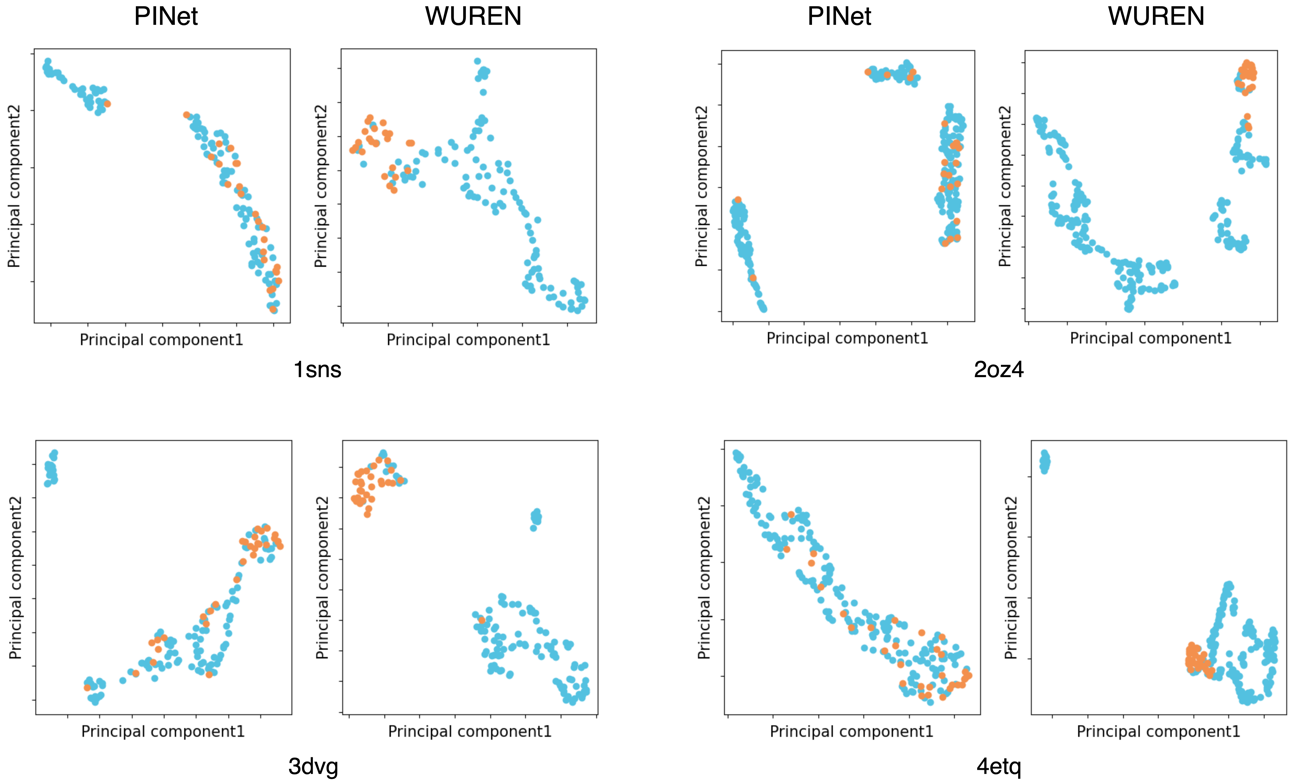

Supplementary Fig. 1: More examples of latent variable spaces. The scatter plots of the penultimate layer’s embedding after PCA, orange represents positive samples, blue represents negative samples.

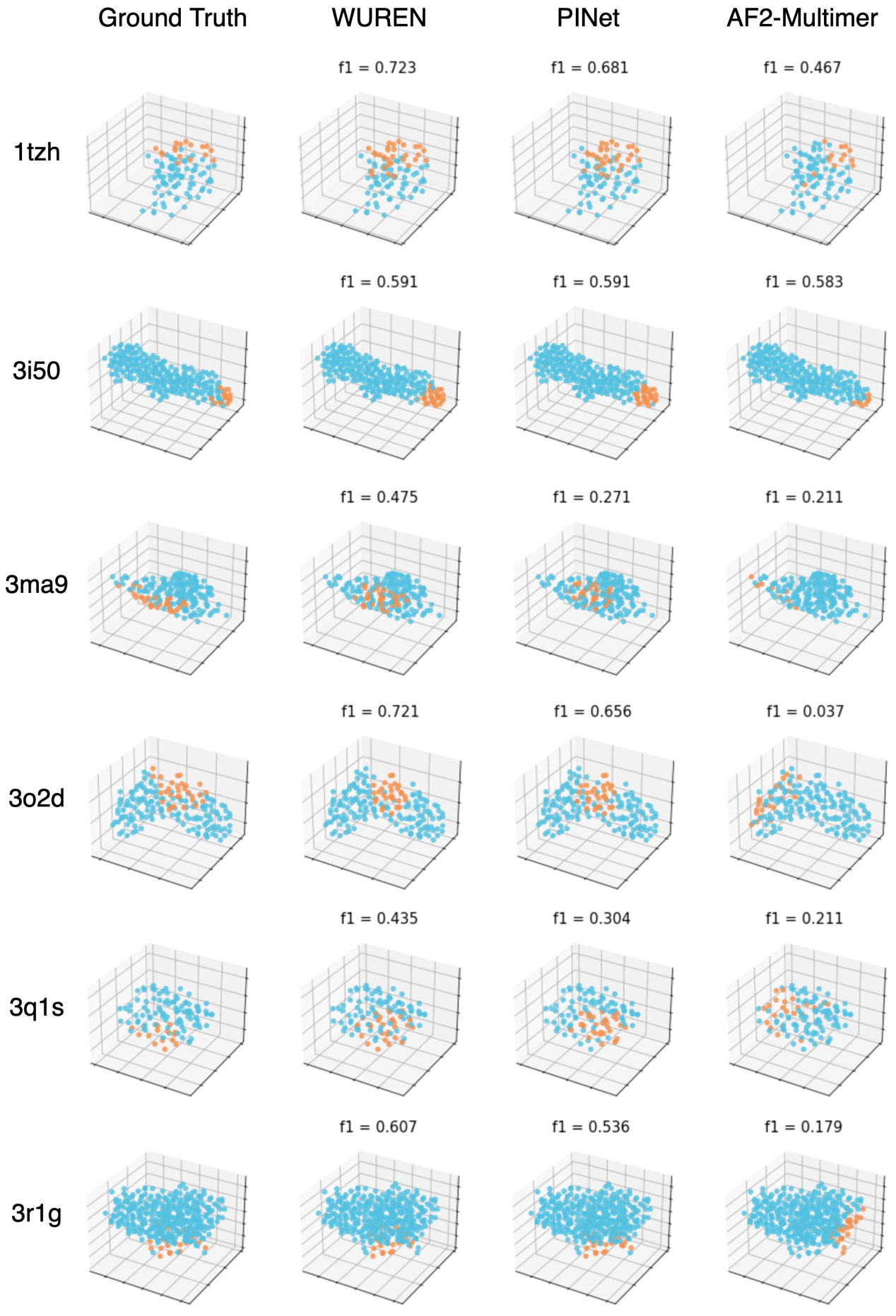

Supplementary Fig. 2: More examples of 3D visualization of prediction results. 3D scatters plots of prediction results of different models, orange represents positive samples, blue represents negative samples. WUREN outperforms PINet and AF2-multimer in general.

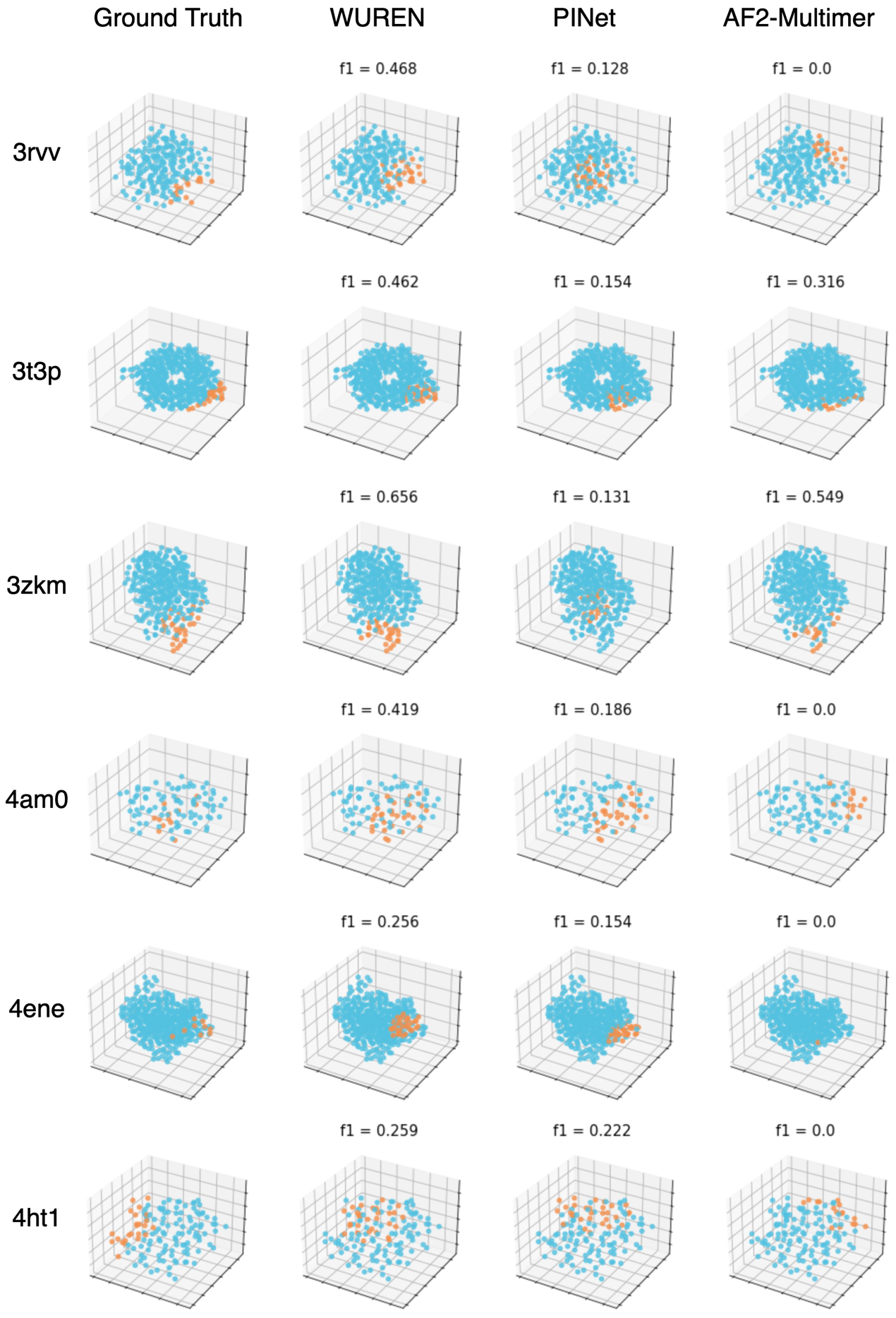

Supplementary Fig. 3: More examples of 3D visualization of prediction results. 3D scatters plots of prediction results of different models, orange represents positive samples, blue represents negative samples. WUREN outperforms PINet and AF2-multimer in general.

### Performance on SAbDab newly released Dataset

The SAbDab newly released data used in this article is shown in the table below.

**Supplementary Table. 2: List of 77 PDB files that constitute the additional dataset for epitope prediction**

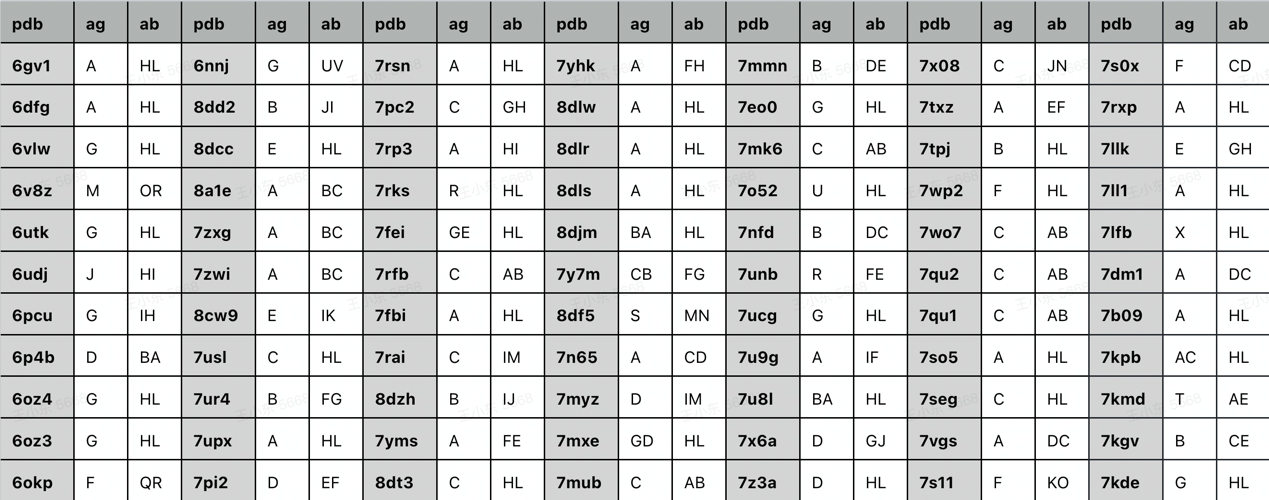

WUREN achieves the best results on this dataset in general.

**Supplementary Table. 3:** Test result table of WUREN on newly published data from SAbDab

| Methods | F1 | Precision | Recall |
| --- | --- | --- | --- |
| xTrimo-Multimer(1*1, unrelaxed) | 0.137 ± 0.03 | 0.141 ± 0.02 | 0.143 ± 0.03 |
| xTrimo-Multimer(5*5, relaxed) | 0.167 ± 0.01 | 0.173 ± 0.01 | 0.175 ± 0.01 |
| AF2-Multimer(1*1, unrelaxed) | 0.172 ± 0.02 | 0.227 ± 0.02 | 0.161 ± 0.02 |
| AF2-Multimer(5*5, relaxed) | 0.231 ± 0.01 | 0.224 ± 0.01 | 0.252 ± 0.01 |
| WUREN | **0.331 ± 0.03** | **0.268 ± 0.02** | **0.577** **± 0.04** |

### Performance on MaSIF Dataset

WUREN achieves the best performance on MaSIF dataset, with AUC-PR 0.516 and AUC-ROC 0.891.

**Supplementary Table. 4:** Test result table of WUREN on MaSIF datasets

| Methods | AUC-PR | AUC-ROC |
| --- | --- | --- |
| SPPIDER | Na | 0.657 |
| MaSIF | Na | 0.874 |
| PINet | 0.454 ± 0.04 | 0.883 ± 0.05 |
| WUREN | **0.516 ± 0.05** | **0.891 ± 0.03** |

### Results of Model Ablation Test

The performance is improved as the complexity of the model increases when more representations are fused together.s

**Supplementary Table. 5:** Comparison table of model ablation experimental test results

| Methods | AUC-PR | AUC-ROC | Precision | Recall | F1 |
| --- | --- | --- | --- | --- | --- |
| Epipred | Na | Na | 0.136 | 0.436 | Na |
| PECAN | 0.226 | 0.655 | 0.157 | 0.730 | Na |
| PINet | 0.368 | 0.687 | 0.216 | **0.774** | Na |
| Cross Attetion PointNet ++ (CAP) | 0.324  (-29.87%) | 0.731  (-16.65%) | 0.321  (-10.58%) | 0.407  (-37.09%) | 0.359  (-22.13%) |
| Cross Attention GCN (CAG) | 0.333  (-27.92%) | 0.778  (-11.29%) | 0.201  (-44.01%) | *0.696*  *(+7.57%)* | 0.311  (-32.54%) |
| Cross Attention Transformer (CAT) | 0.312  (-32.47%) | 0.776  (-11.52%) | 0.188  (-47.63) | *0.649*  *(+0.31%)* | 0.291  (-36.88%) |
| CAP + CAG | 0.341  (-26.19%) | 0.824  (-6.04%) | 0.331  (-7.79%) | 0.587  (-9.27%) | 0.423  (-8.24%) |
| CAP + CAT | 0.372  (-14.52%) | 0.858  (-2.17%) | 0.279  (-22.28%) | *0.754*  *(+16.54%)* | 0.408  (-11.49%) |
| CAG + CAT | 0.386  (-16.45%) | 0.793  (-9.58%) | 0.307  (-14.48%) | 0.642  (-0.77%) | 0.415  (-9.98%) |
| CAP + CAG + CAT (WUREN) | **0.462** | **0.877** | **0.359** | 0.647 | **0.461** |
